# Method to estimate the approximate samples size that yield a certain number of significant GWAS signals in polygenic traits

**DOI:** 10.1101/219733

**Authors:** Silviu-Alin Bacanu, Kenneth S. Kendler

## Abstract

To argue for increased sample collection for disorders without significant findings, researchers retorted to plotting, for multiple traits, the number of significant findings as a function of the sample size. However, for polygenic traits, the prevalence of the disorder confounds the relationship between the number of significant findings and the sample size. To adjust the number of significant findings for prevalence, we develop a method that uses the expected noncentrality of the contrast between liabilities of cases and controls. We empirically find that, when compared to the sample size, this measure is a better predictor of number of significant findings. Even more, we show that the sample size effect on the number of signals is explained by the noncetrality measure. Finally, we provide an R script to estimate the required sample size (non-centrality) needed to yield a pre-specified number of significant findings.

---

To illustrate the tractability of complex diseases, researchers intuitively plot/regress^1,2^ the number of significant findings, n_s_, by the sample size, N, (see Fig. 2 in Kim et al^1^ and Fig. 3 in this paper). Early in the GWAS era such a plot suggested that the number of significant hits is approximately linear after the emergence of the first genome wide significant finding (Mark Daly PGC presentation). While such analyses are definitely informative, for polygenic traits such plots are confounded by the trait prevalences (Fig. 2 in Gratten et all^3^). For a better characterization of trait effect size that is not cryptically influenced by prevalence, we propose an approach to adjust traits for their prevalences and provide an empirical relation between such normalized variables and the number of significant findings for given sample sizes.

Let us assume the existence of biologically informative covariates, e.g. gender and ancestry principal components, which helps us in recovering the liability to disease (even up to a multiplication factor),, for both cases and controls for a binary trait (BT) of prevalence. (It should be noted that working on the liability scale, instead of the natural binary case control scale, is also supported by the Invariance Principle of statistical mathematics^4^, which states that the inference should not be affected by the scale/transformations one chooses to employ.) For the threshold-liability model, let the threshold be *τ_K_ = Φ*^−1^(1 − *K*), where *Φ*^−1^ is the inverse cumulative distribution function of the Gaussian distribution. In a threshold-liability model L ≥ τ_*K*_ for cases and L < *τ_K_* for controls. Thus, for a study consisting of *N*_1_ cases and *N*_2_ controls, the normalized effect size (δ), i.e. the difference in liability between cases and controls after adjusting for its standard error, is:

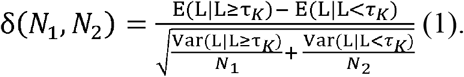

If *φ* is the probability density function for the Gaussian distribution, after substituting the expressions for expectation and variance of truncated Gaussian distributions^5^, relationship (1) becomes:

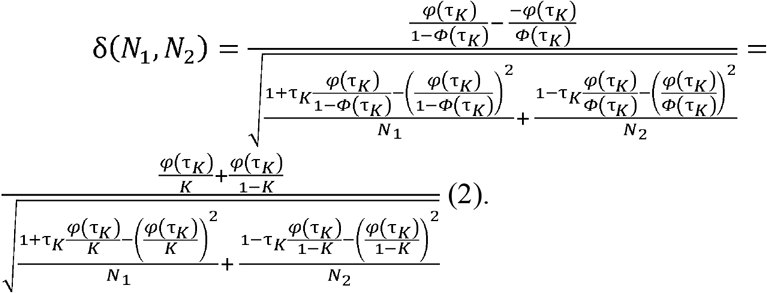

However, most often researchers work with the χ^2^ distribution and, on this scale, the non-centrality parameter of contrasting case and control liabilities is λ(*N*_1_, *N*_2_) = δ^2^(*N*_1_, *N*_2_). In turn, detection power is increasing with increased non-centrality parameter.

While equation (2) is derived for binary traits, it can be extended to quantitative traits (QT). For instance, we can use a first order approximation for QT as a case control trait with prevalence of 50% (i.e. a contrast above median height vs. below median height). While, in practice, such a discretization approach leads to power loss, we stress that the GWAS statistics are already obtained using a QT. The above/below median approximation is only used to extend the use of equation (2). With this preparatory work, the noncentrality per case and control unit (*N*_1_ = *N*_2_ = 1), λ(1,1), increases by ∼60% with a decrease in prevalence (Fig. 1).

**Figure 1.**
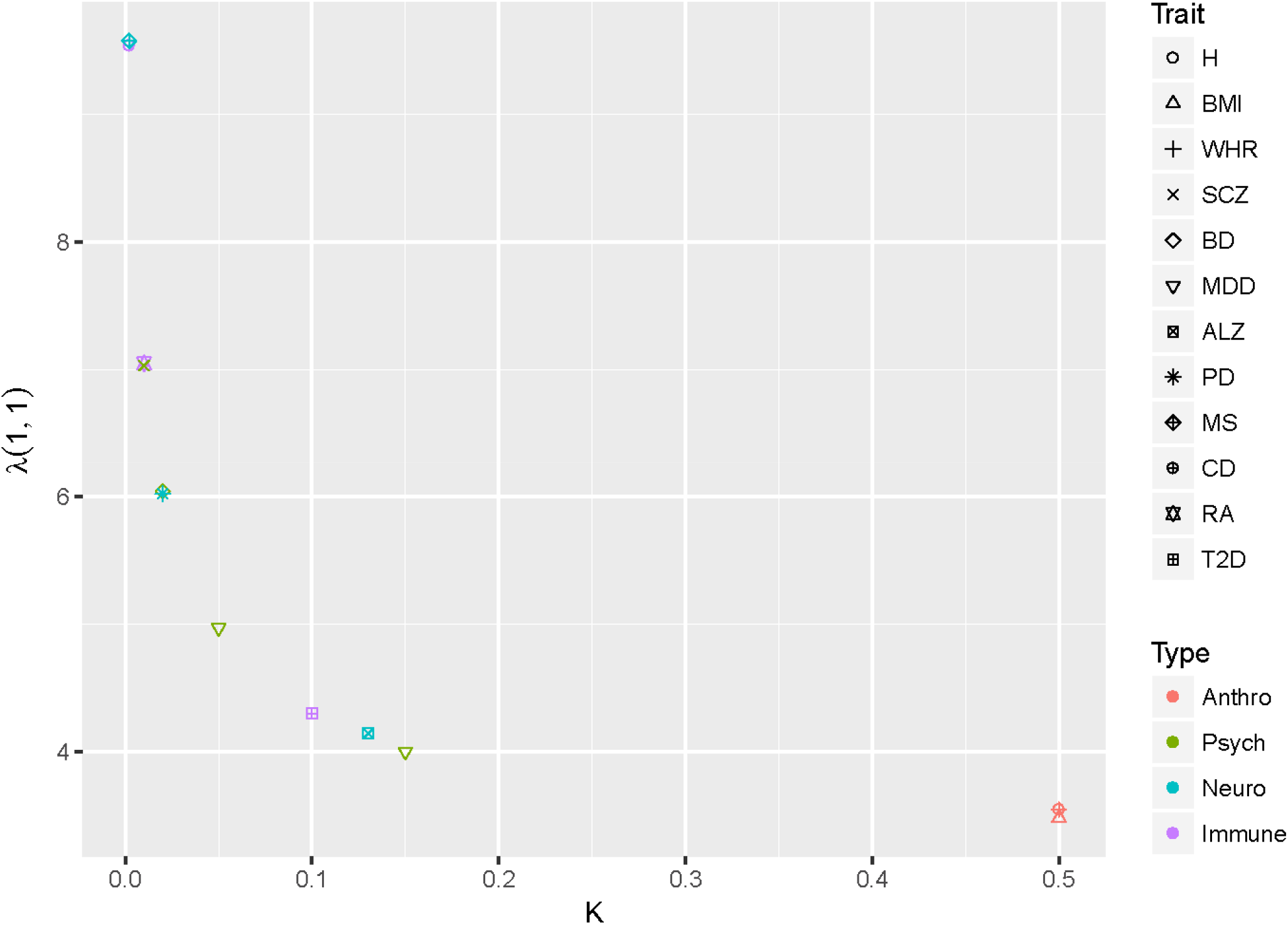
Noncentrality parameter for various traits

To empirically investigate whether λ is a better measure than *N*_1_, or *N* = *N*_1_ + *N*_2_, to describe observed n_s_, we analyze the number of significant findings (Table 1) for multiple studies for some of the most widely investigated traits. Three phenotypes are chosen from each of the four investigated trait classes (see Table 1 for references): anthropometric (all QTs) and psychiatric, neurodegenerative and immune diseases (all BT). Anthropometric traits (denoted as Anthro in plot legends) are height (H), body mass index (BMI) and waist-to-hip ratio (WHR). Psychiatric (Psych) traits are the main psychiatric disorders: schizophrenia (SCZ), bipolar disorder (BD) and major depressive disorder (MDD). As neurodegenerative (Neuro) we chose Alzheimer’s disease (ALZ), Parkinson’s disease (PD and multiple sclerosis (MS). Finally, we chose as immune (Immune) disorders: Crohn’s disease (CD), rheumatoid arthritis (RA) and type 2 diabetes (T2D).

**Table 1.** Table of Studies

| Trait | Abbrev. | K | N cases | N controls | Ns | First author and reference |
| --- | --- | --- | --- | --- | --- | --- |
|  |  |  | 6,800 | 6,800 | 20 | Weedon<sup>6</sup> |
| Height | H | 0.5 | 90,000 | 90,000 | 180 | Lango<sup>7</sup> |
|  |  |  | 125,000 | 125,000 | 423 | Wood<sup>8</sup> |
|  |  |  | 2,500 | 2,500 | 1 | Scuteri<sup>9</sup> |
|  |  |  | 16,000 | 16,000 | 8 | Willer<sup>10</sup> |
| Body Mass Index | BMI | 0.5 | 16,500 | 16,500 | 11 | Thorleifsson<sup>11</sup> |
|  |  |  | 125,000 | 125,000 | 32 | Speliotes<sup>12</sup> |
|  |  |  | 170,000 | 170,000 | 97 | Locke<sup>13</sup> |
|  |  |  | 20,000 | 20,000 | 1 | Lindgren<sup>14</sup> |
| Waist to Hip Ratio | WHR | 0.5 | 40,000 | 40,000 | 14 | Heid<sup>15</sup> |
|  |  |  | 110,000 | 110,000 | 49 | Shungin<sup>16</sup> |
|  |  |  | 17,500 | 33,500 | 7 | PGC1<sup>17</sup> |
| Schizophrenia | SCZ | 0.01<sup>18</sup> | 20,000 | 37,000 | 22 | PGC1.5<sup>19</sup> |
|  |  |  | 37,000 | 113,000 | 108 | PGC2<sup>20</sup> |
|  |  |  | 2,000 | 3,000 | 1 | WTCCC<sup>21</sup> |
| Bipolar Disorder | BD | 0.02<sup>22</sup> | 7,500 | 9,250 | 2 | PGC1<sup>23</sup> |
|  |  |  | 10,000 | 15,000 | 5 | Muhleisen (personal communication) |
| Major Depressive Disorder | MDD | 0.15<sup>24</sup> | 9,000 | 9,500 | 0 | PGC MDD<sup>25</sup> |
| (Recurrent MDD) |  | (0.05)<sup>26</sup> | 6,000 | 6,000 | 2 | CONVERGE<sup>26</sup> |
|  |  |  | 131,000 | 330,000 | 45 | PGC2 MDD (online presentation) |
|  |  |  | 8,300 | 7,300 | 3 | Naj<sup>27</sup> |
| Alzheimer's Disease | ALZ | 0.13<sup>27</sup> | 17,000 | 37,000 | 15 | Lambert<sup>28</sup> |
|  |  |  | 25,000 | 48,000 | 20 | Lambert<sup>28</sup> |
|  |  |  | 1,700 | 4,000 | 2 | Simon-Sanchez<sup>29</sup> |
| Parkinson's Disease | PD | 0.02<sup>30</sup> | 3,500 | 30,000 | 8 | Do<sup>31</sup> |
|  |  |  | 13,700 | 95,000 | 26 | Nalls<sup>30</sup> |
|  |  |  | 1,000 | 900 | 1 | Baranzini<sup>32</sup> |
| Multiple Sclerosis | MS | 0.002<sup>32</sup> | 4,800 | 9,300 | 6 | De Jager<sup>33</sup> |
|  |  |  | 9,800 | 17,400 | 23 | IMSGC<sup>34</sup> |
|  |  |  | 1,000 | 2,345 | 12 | McGovern<sup>35</sup> |
| Crohn's Disease | CD<sup>35</sup> | 0.002 | 3,250 | 4,800 | 33 | Barrett<sup>36</sup> |
|  |  |  | 6,350 | 15,050 | 71 | Franke<sup>37</sup> |
|  |  |  | 2,100 | 2,500 | 4 | Jiang<sup>38</sup> |
| Rheumatoid Arthritis | RA | 0.01<sup>38</sup> | 5,500 | 20200 | 10 | Stahl<sup>39</sup> |
|  |  |  | 20,000 | 620,00 | 57 | Okada<sup>40</sup> |
|  |  |  | 661 | 614 | 5 | Sladek<sup>41</sup> |
| Type 2 Diabetes | T2D | 0.1<sup>41</sup> | 1,464 | 1,467 | 5 | Diabetes at BROAD<sup>42</sup> |
|  |  |  | 5,500 | 14,500 | 20 | Kooner<sup>43</sup> |
|  |  |  | 26,500 | 84,000 | 26 | DIAGRAM<sup>44</sup> |

To assist in predicting n_s_ as a function of λ, we also need to determine what transformation should we use for n_s_ and λ/*N*_1_ to make the relationship between n_s_ and λ stronger. As mentioned in the introduction, the intuitive idea is to use the identity scale, i.e. no transformation. However, given that n_s_ can be viewed as a sum of Bernoulli variables (0-non-significant and 1 significant), Chernoff inequality^4^ suggests that a log transformation of n_s_ is likely much more desirable. For effect sizes λ (and likely, as its transformation, *N*), the plotting of the log probability of a significant signal (*α* = 5*x*10^−8^) as a function of noncentrality, λ, and its log transformation, also show a much better fit (Fig. 2) for the log transformation (R^2^ of 99.4% vs 91%). Given that the probability of a significant find is proportional to the number of significant findings, this suggests that the log transformation is also suitable for λ.

**Figure 2.**
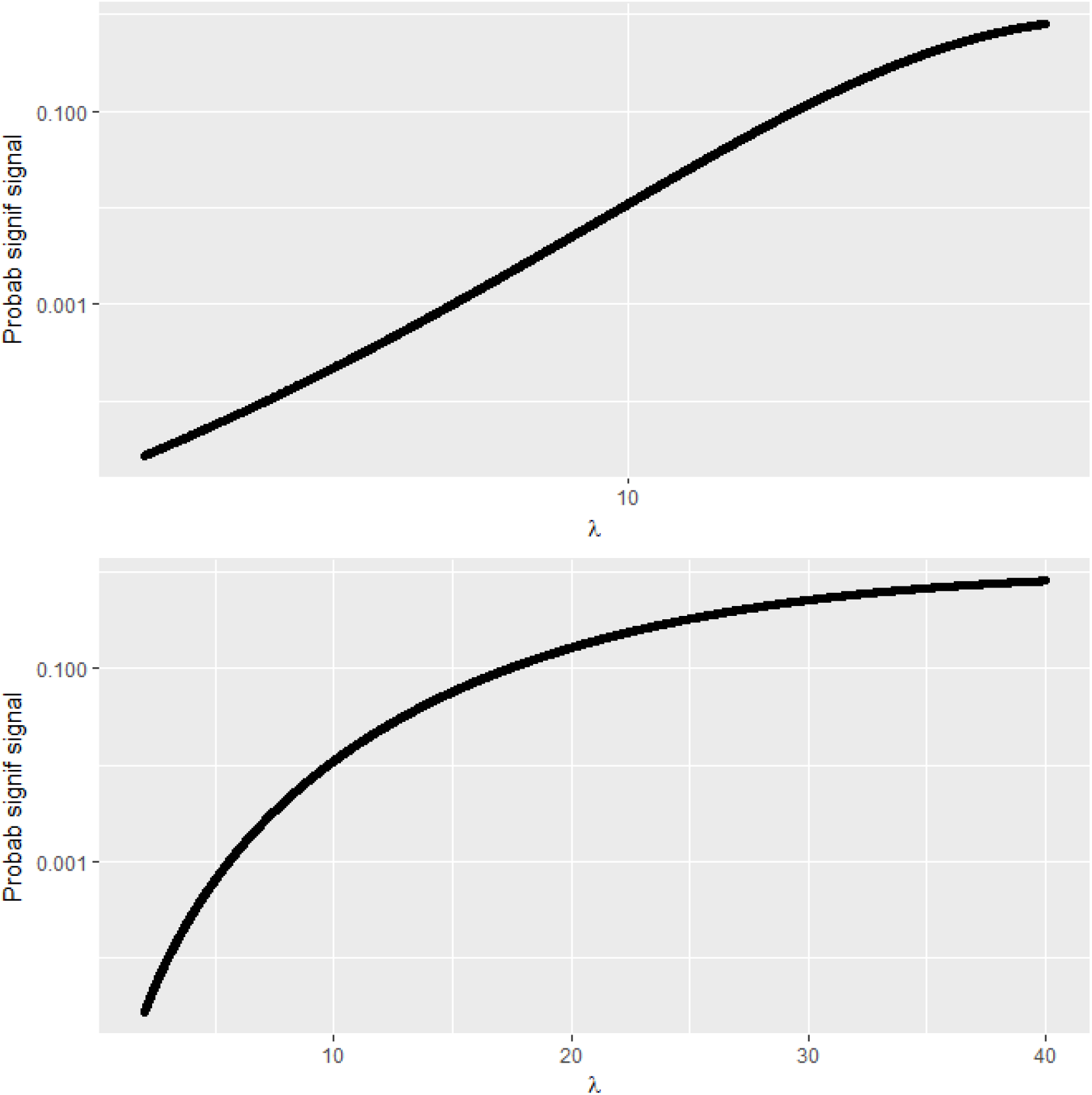
The probability of a significant signal (log scale) as a function of noncentrality on log scale (above) and identity scale (below).

Thus to establish the relationship between regressing log[n_s_] and log(λ) (also log *N*_1_, log *N*) we use a gls model (in nlme R package) assuming an autoregressive of order 1 (AR1) correlation structure for observations within the same trait (due to earlier studies being included in all subsequent meta-analysis of this disease). We used the model to test whether the effects of *N* and *N*_1_ on n_s_ are mediated only via log(λ), i.e. we regressed log[n_s_] on log(λ), log[N], log(*N*_1_), N and λ_1_. In this model, only log(λ) was significant (p-value of 0.025) and all the others were not (p-values of 0.58 and 0.73). Even more, stepwise elimination on non-significant variables left only log(λ) as significant with log[N] being the last to be eliminated with a p-values of 0.65. This result strongly suggests that the effect of N and *N*_1_ on n_s_ is wholly mediated by λ and thus non-centrality is a better predictor than sample size. The gls model was also used to vividly illustrate the better performance of our theoretically chosen transformations: when using the natural sample size scale for both n_s_ and (Fig. 3), the fit (R^2^=0.42) is much poorer than using log scale for both (Fig. 4) (R^2^=0.71). (The similar in spirit square root transformation of λ performed only moderately worse than log.)

**Figure 3.**
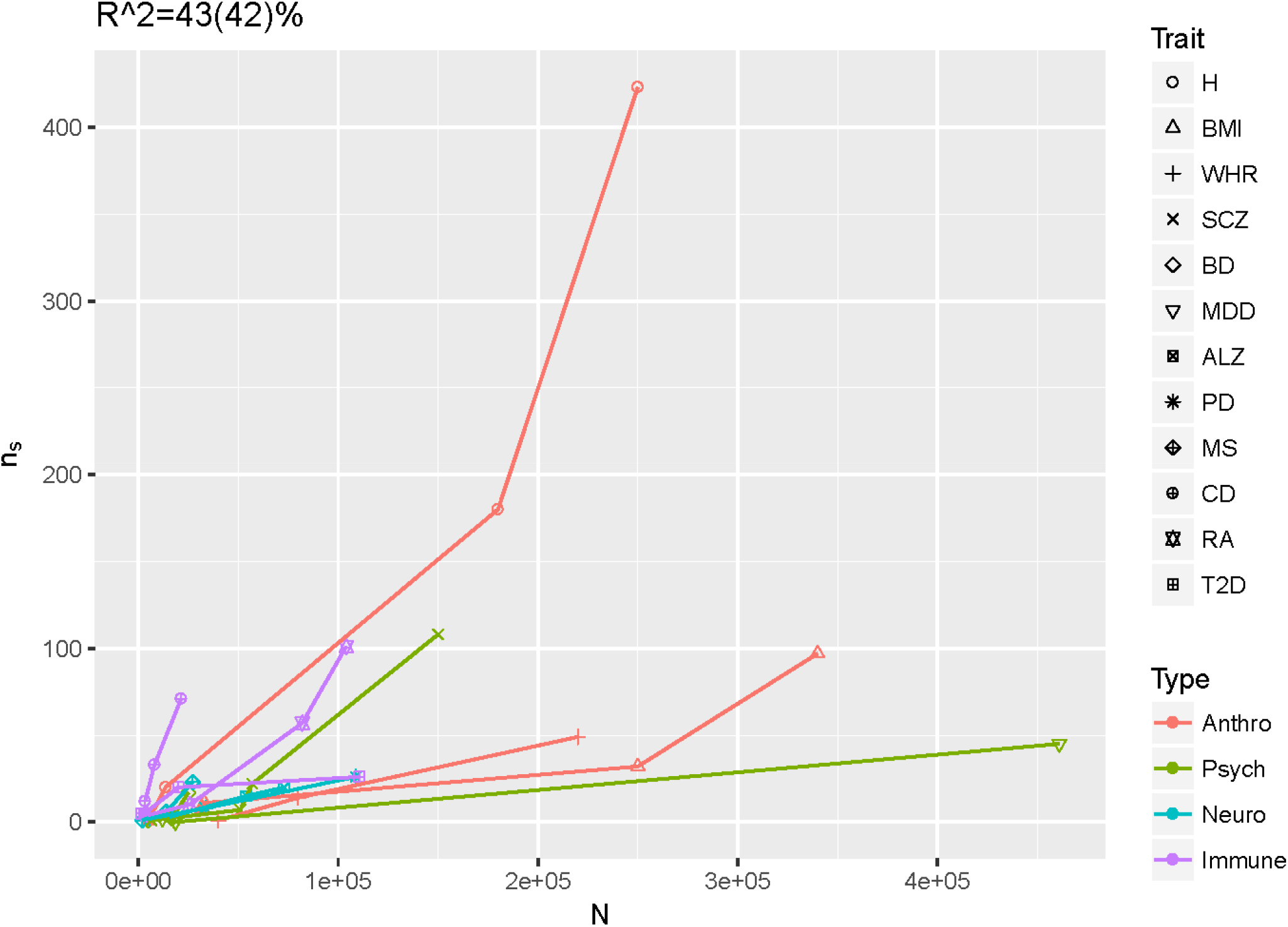
Number of significant findings vs. sample size (without type 2 diabetes-T2D).

**Figure 4.**
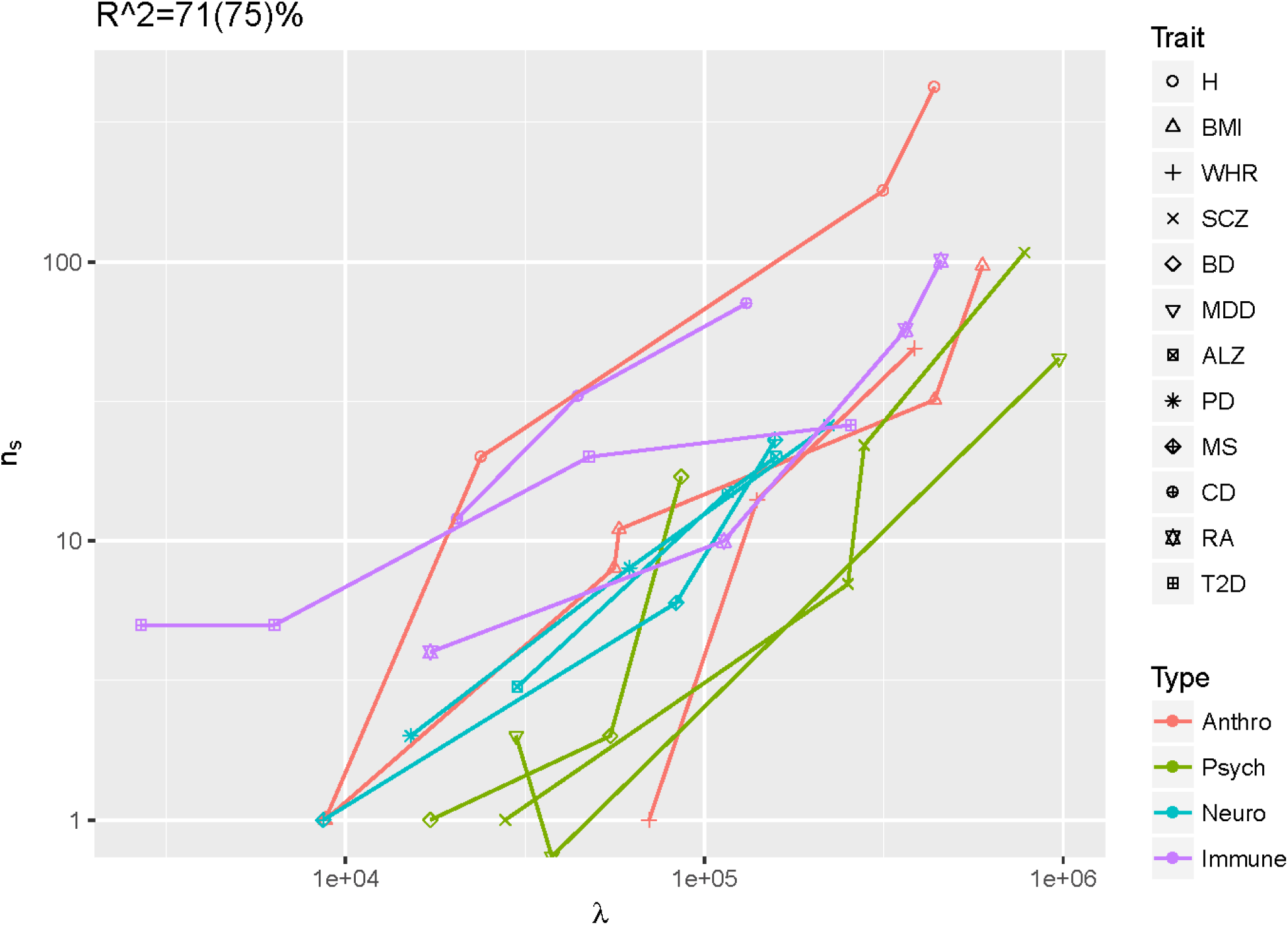
Number of significant findings vs noncentrality parameter (without T2D). Both axes are log scale.

We stress again that the above results suggest that our proposed measure on log scale better predicts the (log) number of significant findings for traits of various prevalences. Thus, λ from relationship (1) is a desirable effect size measure that is not confounded by prevalence. Based on the gls regression of log[n_s_] on log(λ), the best prediction for the number of significant findings is:

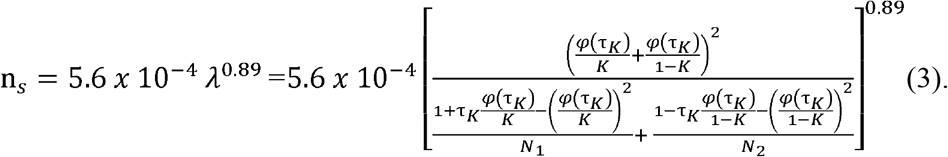

However, most of the time the researchers want to estimate the number of cases, *N*_1_, needed to obtain a certain number of significant findings, n_*s*_. To this end let *N*_2_ = *q N*_1_, where generally *q* > 1 is largely known. Then equality (3) can be solved for *N*_1_, as follows:

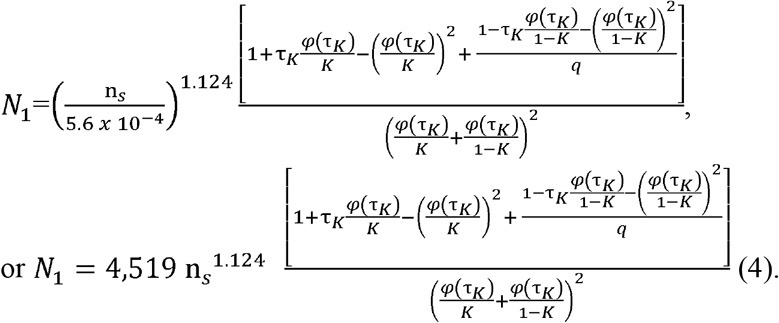

To assist applied researchers in sizing their studies, we present in the Appendix the R implementation of equalities (3) and (4).

## Appendix

~~~
**# R function for estimating the noncentrality and # cases
*## K-prevalence, Nca - # cases ; Nco - # controls***
get.nonc<-function(K=K, Nca=Nca, Nco=Nco){
      tau.K<-qnorm(K, low=F)
      ncp<-(dnorm(tau.K)*(1/K+1/(1-K)))^2/((1+tau.K*dnorm(tau.K)/K-(dnorm(tau.K)/K)^2)/Nca+(1-tau.K*dnorm(tau.K)/(1-K)-(dnorm(tau.K)/(1-K))^2)/Nco)
      ncp
}
~~~

~~~
**# R function for estimating the required # cases yielding # of signals using our formula (4)
*## K-prevalence, ns - # desired significant findings & q=ratio of controls to cases (often q>1)***
get.n.cases<-function(K=K, ns=1, q=1){
     tau.K<-qnorm(K, low=F)
     N1<-4519*ns^1.124*((1+tau.K*dnorm(tau.K)/K-(dnorm(tau.K)/K)^2)+(1-tau.K*dnorm(tau.K)/(1-K)-(dnorm(tau.K)/(1-K))^2)/q)/(dnorm(tau.K)*(1/K+1/(1-K)))^2
     N1
}
~~~

